# Homologue Series Detection and Management in LC-MS data with *homologueDiscoverer*

**DOI:** 10.1101/2022.07.20.500749

**Authors:** Kevin Mildau, Justin J.J. van der Hooft, Mira Flasch, Benedikt Warth, Yasin El Abiead, Gunda Koellensperger, Jürgen Zanghellini, Christoph Büschl

## Abstract

**Summary:** Untargeted metabolomics data analysis is highly labor intensive and can be severely frustrated by both experimental noise and redundant features. Homologous polymer series are a particular case of features that can either represent large numbers of noise features, or alternatively represent features of interest with large peak redundancy. Here we present *homologueDiscoverer*, an R package which allows for the targeted and untargeted detection of homologue series as well as their evaluation and management using interactive plots and simple local database functionalities.

**Availability:** *homologueDiscoverer* is freely available at github https://github.com/kevinmildau/homologueDiscoverer.

## 1 Introduction

Untargeted metabolomics techniques based on LC-MS/MS allow for the generation of comprehensive snapshots of the chemical composition of samples and find wide use in diverse fields ranging from clinical biomarker discovery to natural product discovery (Kennedy *et al*., 2018; Tsugawa *et al*., 2021). However, LC-MS data are plagued by bioinformatic and chemical noise as well as redundancies which can hamper data analysis steps such as metabolite annotation or molecular networking (Schiffman *et al*., 2019; da Silva *et al*., 2018). One common type of contamination is caused by homologue series, i.e., groups of compounds with differing numbers of the same repeating unit. Existing tools for dealing with homologue series either exclusively work with specific pre-specified increments (da Silva *et al*., 2018) or tend to produce highly complex outputs that make manual curation to discriminate biologically meaningful series very difficult (Loos and Singer, 2017).

In this work we present *homologueDiscoverer* (*HD*), an R package which allows for the detection and processing of homologues within LC-MS/MS peak tables. Homologues are frequently encountered in LC-MS/MS runs and tend to exhibit highly regular MS1 patterns (da Silva *et al*., 2018; Loos and Singer, 2017). Indeed, groups of peaks stemming from the same homologue polymer but with different numbers of the repetitive unit exhibit nearly identical mass-to-charge ratio steps between peaks while also showing systematic trends in retention time. *homologueDiscoverer* capitalizes on these systematic trends by extracting homologue series using either pre-specified increments or untargeted search windows. Homologue detection allows reducing data complexity through meaningful feature grouping and provision of feature sets for exclusion from further analysis, thereby allowing researchers to focus on biologically relevant information.

## 2 Application Features

*homologueDiscoverer* provides pre-specified step size (targeted) and untargeted homologue detection algorithms. In both cases, the algorithm iterates through the mz and rt sorted peak table and deploys a two phase series detection routine from each peak. In the initialization phase of the algorithm it is checked whether a second peak lies within the search window of the algorithm relative to the root peak. In the targeted search mode, specific mass-to-charge ratio increments or decrements in conjunction with retention time constraints lead to the establishment of very few 2-tuples. In contrast, the untargeted mode creates long lists of 2-tuples of peaks that are a combination of the current root peak and any peak within the specified search window. Once initialized, the algorithm proceeds to the second phase of series extension, where for each 2-tuple created in the first phase, the corresponding mass-to-charge ratio increment and retention time constraints are used to screen the peak table for any potential candidates for series extension. At three series candidates or more the algorithm deploys further heuristics to restrict retention time windows. Specifically, the retention time step trend of the first 3 series members is used to assess whether retention time steps are either independent of mass, or respectively increasing or decreasing with mass. These trends are used to constrain search windows accordingly. Candidate series are grown until no further candidates match the constraints imposed, and the longest series exceeding the minimum series length is selected as a homologue series (more detailed description in *Supplementary Information*). Our greedy algorithm set-up extracts any peaks grouped to belong to a homologue series from the peak table before continuing its search, thereby guaranteeing that each peak will be annotated to belong to only one homologue series. This feature leads to a very concise and readable output which can be evaluated in interactive shiny graphs (Figure 1) or stored in augmented peak tables for use in homologue annotation of new samples using functions provided by *homologueDiscoverer*.

**Fig. 1.**
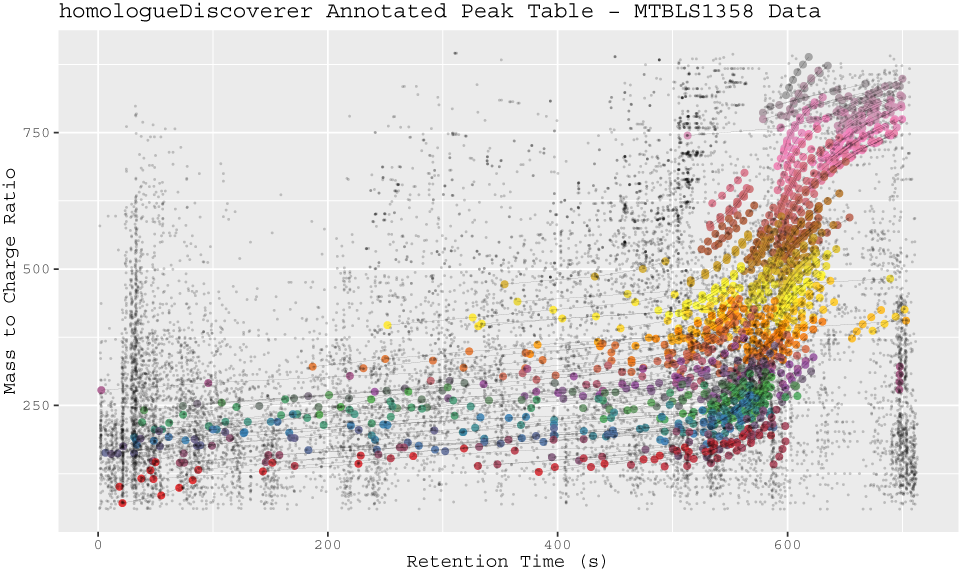
Annotated peak table for untargeted homologue detection run on MTBLS1358 dataset. Homologue series are connected by lines and their peaks are highlighted using larger point sizes. Despite the narrow search settings used, more than 8 percent of peaks were grouped into homologue series, highlighting data reduction potential.

## 3 Results & Discussion

We have evaluated our tool on the PEG17 and PEG70 polyethylene glycol spiked plasma datasets of da Silva *et al*. (2018). While many of the peaks in the samples can be attributed to polymers, not all polymers exhibit the type of characteristic MS1 homologue data trends our algorithm is based on. In our evaluation runs, 115 of 276 PEG17 peaks and 140 of 251 PEG70 peaks were annotated to belong to homologue series. Hence, roughly half of the polymer peaks can be grouped via characteristic trends. Moreover, 7 and 10 peaks not originating from the spiked polymers were grouped into homologue series for PEG17 and PEG70 spiked samples, respectively (see *suppl. section 2*). When applying a comparatively narrow search window run of *homologueDiscoverer* (*HD*) to the larger human cell model MTBLS1358 dataset of Flasch *et al*. (2020) we find 275 homologue series groupings consisting of 1372 peaks providing a substantial amount of feature grouping (see *suppl. section 3*). In addition, we show that *HD* manages to find both homologues series whose mass-to-charge ratio increases and decreases over retention time when applied to yeast extract data (see *suppl. section 4*).

We further compared our tool to *nontarget* (*NT*) (Loos and Singer, 2017) on PEG spiked samples as well as on the MTBLS1258 dataset. Homologue detection comparisons on PEG spiked data show large discrepancies in exact series overlap (exactly identical peak members in identical order), but large overlaps in feature peaks grouped into homologue series (see *suppl. section 2*). Comparison runs between *HD* and *NT* on the MTBLS1358 dataset showed larger discrepancies in both grouped peaks and exact series correspondence (see *suppl. section 3*). Hence, both tools find very similar sets of peaks to be part of series in the simple PEG spiked datasets, although discrepancies on the nature of the series can be large. For more complex datasets such as MTBLS1358, differences between *HD* and *NT* become larger, with results being highly sensitive to settings used. The algorithms are not directly comparable. However, *NT* allows for the same peak to be contained in several series, while *HD* is constructed such that each peak can only be part of one homologue grouping. In addition, constraints built into series extension of *HD* may be more stringent at low series lengths. While both approaches have their merits, we argue that the increased stringency and avoidance of overlaps by *HD* leads to more coherent and human readable output.

## 4 Conclusion

*homologueDiscoverer* makes the detection and management of homologues easier, providing a complete suite of detection algorithms, interactive visualizations, and storage functions. We expect that *homologueDiscoverer* will benefit researchers working with samples which are routinely polluted with homologue series or who wish to study biologically relevant homologue series in their data. This will be especially useful when homologue series cannot be removed from data via blank subtraction or when they present a biologically meaningful features.

## Acknowledgements

The authors thank the members of the Mass Spectrometry Center at the University of Vienna.

## Supplementary information

Supplementary data are available at *Bioinformatics* online.

