## Supplementary Materials for "Homologue Series Detection and Management in LC-MS data with *homologueDiscoverer*"

**5 Bibliography** **22**

### 1 homologueDiscoverer Algorithm Description

*homologueDiscoverer* makes use of a recursive filtering approach to determine whether homologue series are contained within the peak table. The peak table must contain mass-to-charge ratio, retention time and intensity (for later post-processing) value columns. The untargeted search for homologue series extraction requires the following parameters: mz_min and mz_max as well as rt_min and rt_max to define the initial search window for the chain initialization step described later. In addition, a match tolerance in parts per million(ppm) needs to be provided for mass-to-charge ration variability tolerance. In addition to retention time windows, wide and narrow relative retention time tolerance parameters for the expandChain candidates step are required (described in more detail below).

For the untargeted approach, the main algorithm steps are the following.

1. Pre-process peak table to contain mass-to-charge ratio (mz), retention time (rt), intensity (intensity), and peak identifier (peak_id) columns.
2. Sort the peak table by mz and retention time. Ascending or descending order is used depending on whether search is looking for incrementing or decrementing mass-to-charge ratio over retention time.
3. While there are peaks in the sorted peak table do the following:
    (a) Select the first peak in the peak table as the current root peak.
    (b) *Initialize Chain Candidates*: Assess whether there is a peak within the mass-to-charge ratio and retention time window relative to the current root peak (Figure 1 A). Create a list of chain candidate tuples consisting of the root peak and a single possible follow up peak each.
    (c) If there are chain candidates, proceed to the recursive *Expand Chain Candidates* step.
        i. For each chain candidate, use the initialized 2-tuple of peaks to give a precise mass-to-charge ratio step size and a first retention time step size.
        ii. Assess whether there are candidate peaks matching the narrow mz increment and rt step with a wide margin around the rt step size. For each possible match create a new chain of candidates with it as the last set member (Figure 1 B).
        iii. If chains were elongated, proceed to repeat elongation of chain candidates with an additional relative retention time constraint based on the first two retention time step sizes between the first two peak members of the current investigated chain (Figure 1 C). If rt step size is decreasing, only allow for a wide relative margin around the rt step size into the decreasing direction. If rt step size is increasing, only allow for the wide relative margin in the increasing direction. Allow for a narrow relative margin in the opposing direction of the rt trend to account for static retention time step sizes.
        iv. Repeat series chain elongation step iii until all possible series branches have been grown to their maximum with no further matches for elongation.
    (d) Evaluate whether any series exceeds the minimum series length (3 series members would be the absolute minimum number). If multiple series do, select the longest series (Figure 1 D).
    (e) If a valid series has been found, remove all its constituent peaks from the peak table with a common identifier. If no valid series has been found, remove the current root peak from the peak table.
4. Return the annotated peak table with homologue series identifiers.

The algorithm thus proceeds through the peak table in an ordered fashion, greedily selecting any peak series matching the search parameter constraints. Targeted runs are virtually identical to untargeted runs except for in the initialization of chain candidates. Here, the targeted approach deploys a very narrow mass-to-charge ratio increment rather than a wide window, drastically limiting the number of possible candidate chains. The retention time window approach remains identical. Homologue series stored within series data base augmented peak tables can be used for targeted annotation runs including narrow tolerance on both mass-to-charge ratio and retention time.

The first best selection approach of *homologueDiscoverer* leads to concise homologue series database output by avoiding any multiple series assignments to the same peaks. However, it may miss certain valid configurations of peaks that could be considered homologues under different search settings. This means that runs with different settings of homologueDiscoverer may find conflicting annotations for the same peaks. A chemically relevant example of this would be polymers with two different repeating units, where each unit may be found to be part of a different homologue series. However, extraction of constituent peaks in a greedy fashion may select the first valid series and may omit the second possible homologue series since necessary peaks were removed from the search set.

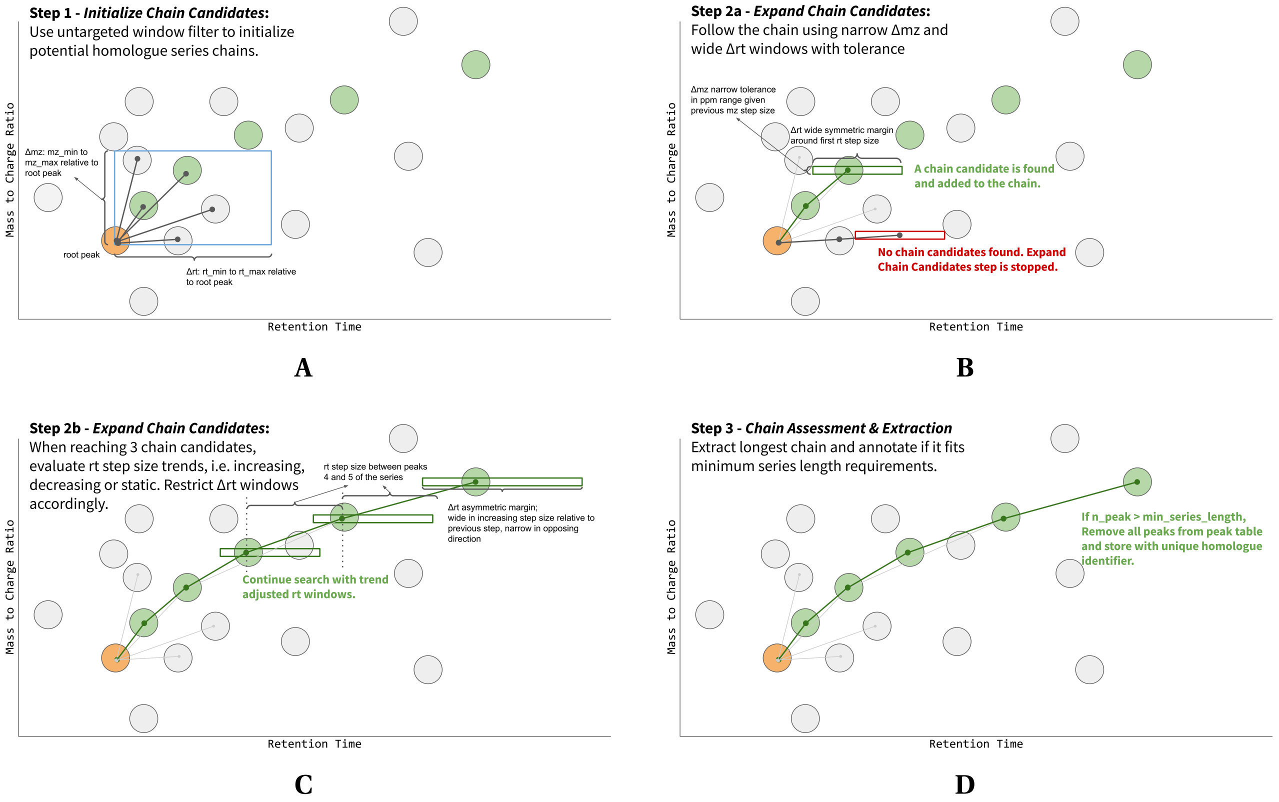

Figure 1: Figures illustrating the main steps of the heuristic chain detection algorithm. **A**) Candidate Chain Initialization using the untargeted window method. Every peak within the search window is used to create a 2-tuple representing the first to peaks of a possible homologue series. **B**) In stage a) of the expand chain candidates step the algorithm uses narrow mz windows and wide symmetric retention time windows to find a possible third member of the chain. **C**) In stage b) of expand chain candidates detected trends in retention time step size over mass-to-charge ratio in the first three peaks of the chain are used to restrict retention time windows in accordance with trends. For instance, if retention time step size tends to increase, decreasing retention time step sizes are not accepted. **D**) Once all possible chains for a specific root node are fully explored, the longest chain is compared against minimum series length requirements. If it exceeds the minimum length, all corresponding peaks are extracted from the peak table and assigned a common homologue series identifier. Screening windows are not to scale. In practice the mass-to-charge ratio range will be within a few ppm of the target mass to charge ratio, while retention time windows will have relatively large margins of the previous step size +− about .5 this same step size into either the positive or negative direction depending on previous trends.

#### 2 PEG Data Validations & Comparisons

*homologueDiscoverer* was run on the PEG17 and PEG70 spiked plasma sample peak tables published by da Silva et al. [1] using the untargeted, increment mode. Search windows were set to between 5 and 100 for the mass to charge ratio and 0.1 to 100 seconds for retention time. Minimum series length was set to 4, with a tolerance in ppm of 20. For comparative purposes, *nontarget* was also run on the same data [2]. *nontarget* runs used identical search window settings or default settings, except for rttol which was set to 50 seconds to allow for differences in retention time step sizes between peaks. These settings were found to lead to rather comparable results for both tools, which as a result of algorithm design differences cannot be made to exactly agree. The code used to generate figures is available on github https://github.com/kevinmildau/homologueDiscoverer-validation-and-comparison.

##### 2.1 PEG Data Annotated Peak Tables

Figures 2 and 3 show annotated peak tables produced by *homologueDiscoverer* for the PEG17 and PEG70 spiked plasma samples respectively. Pseudo annotated peak tables for *nontarget* runs are shown in figures 4 and 5 for PEG17 and PEG70 spikes samples respectively.

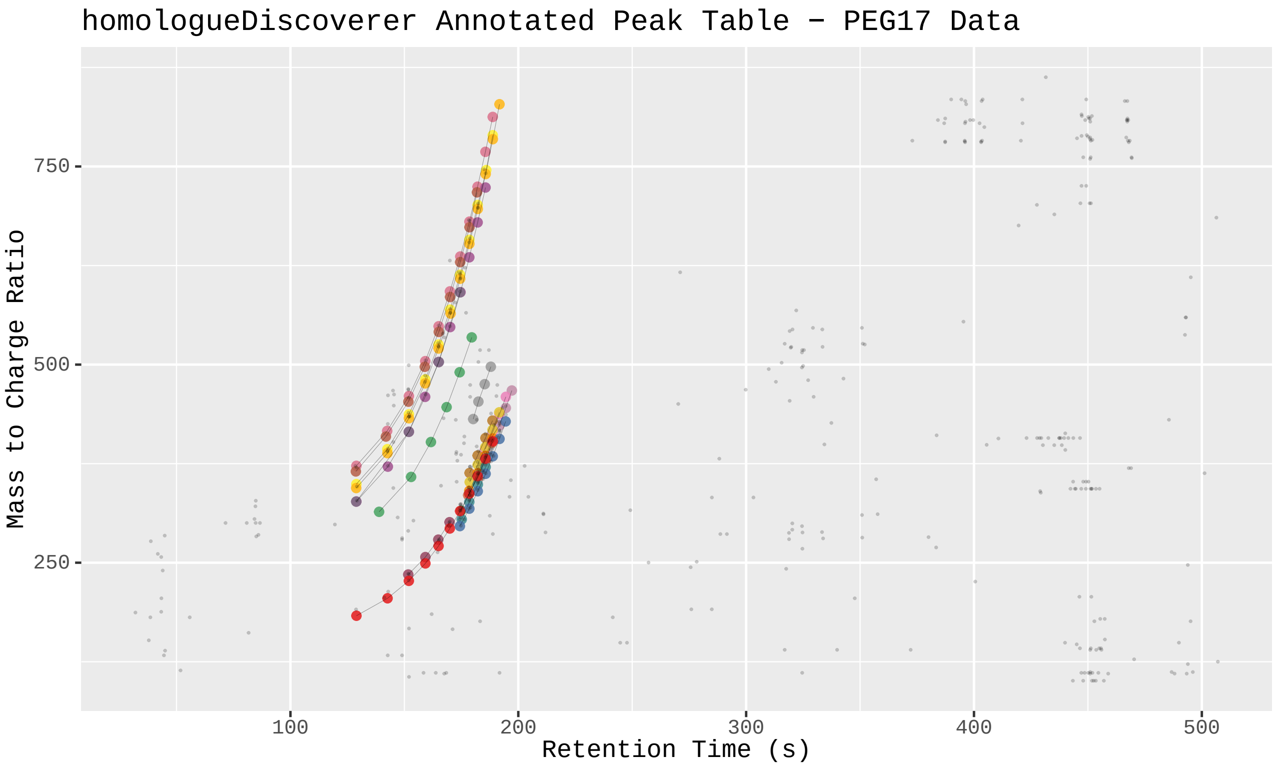

Figure 2: *homologueDiscoverer* output for plasma samples spiked with PEG17. Peaks not annotated to be part of homologue series are shown as faint gray points. Homologue series peaks are emphasized in size. Members of the same homologue series are connected by a line and share the same color.

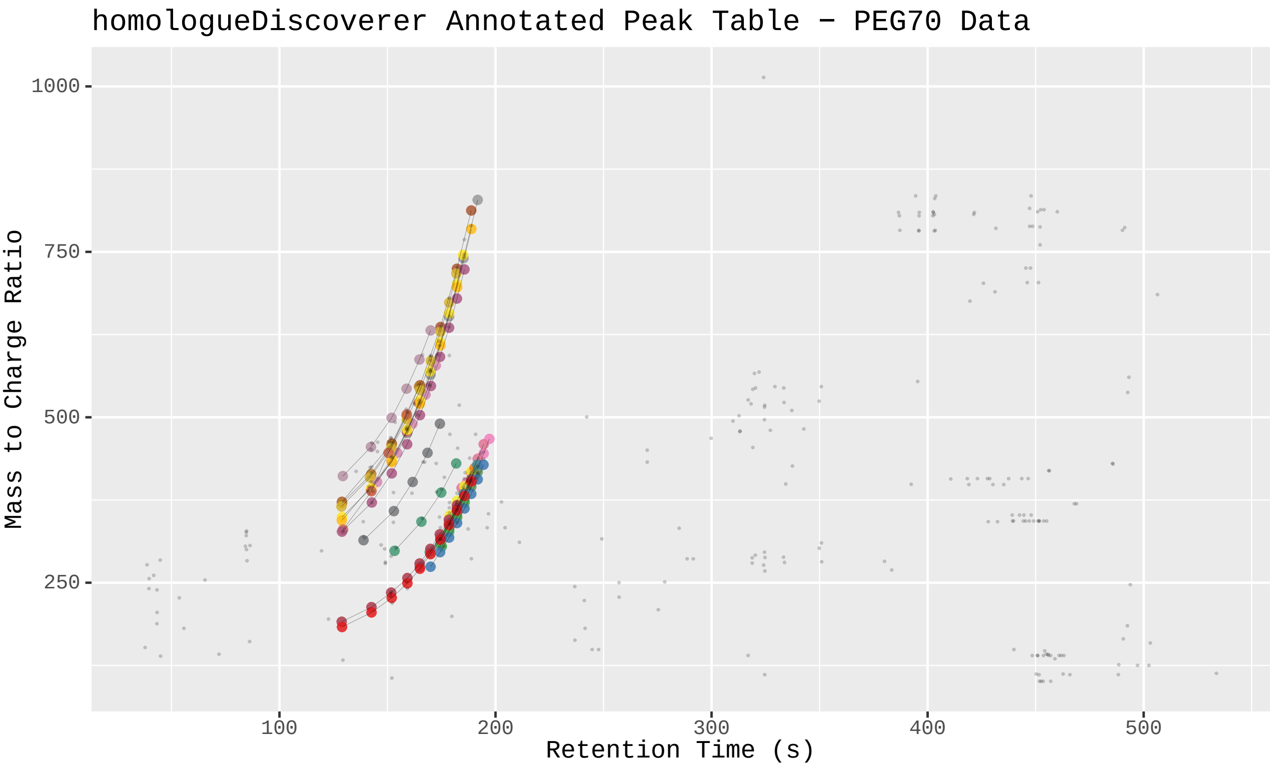

Figure 3: *homologueDiscoverer* output for plasma samples spiked with PEG70. Peaks not annotated to be part of homologue series are shown as faint gray points. Homologue series peaks are emphasized in size. Members of the same homologue series are connected by a line and share the same color.

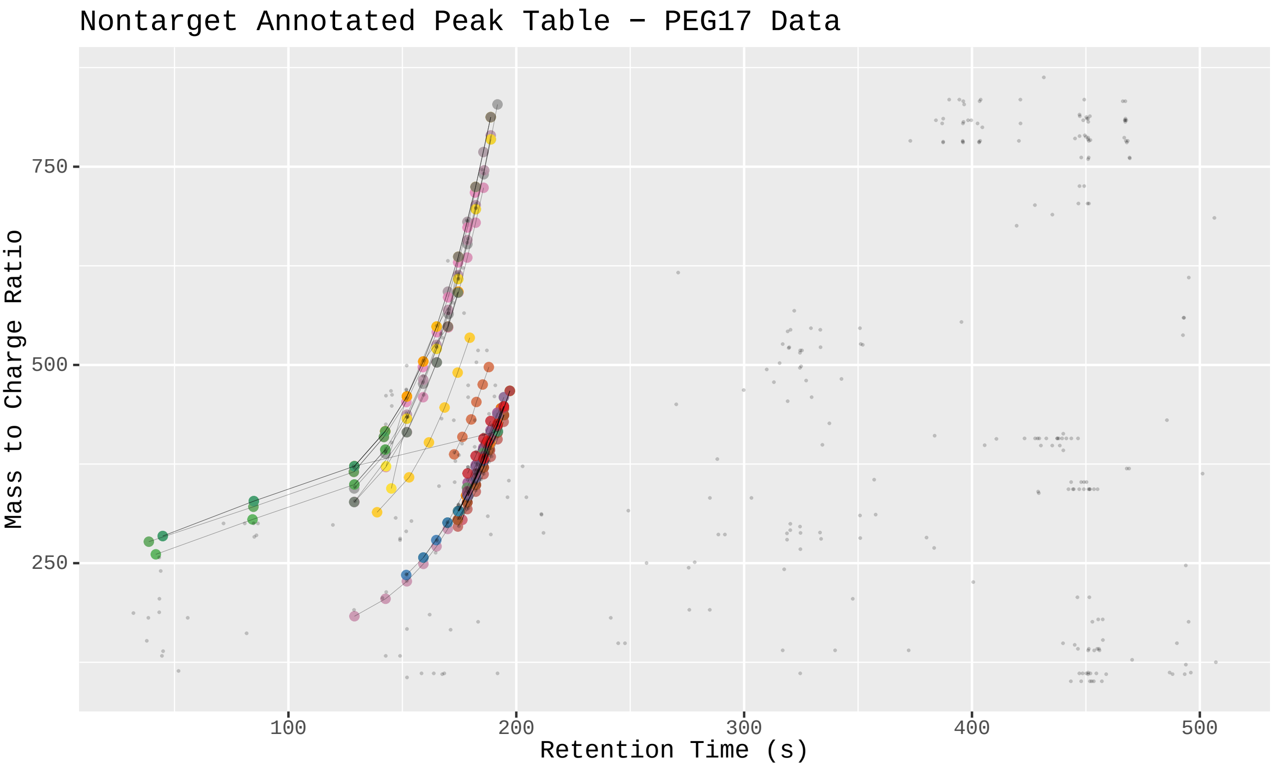

Figure 4: *nontarget* output for plasma samples spiked with PEG17. Peaks not annotated to be part of homologue series are shown as faint gray points. Homologue series peaks are emphasized in size. Members of the same homologue series are connected by a line and share the same color. A custom visualization was implemented to work with *nontarget* output. Peak tables contain duplicate entries for each peak with multiple series assignments. Any peaks highlighted in size that represent homologue series members may thus be overlapping with other points illustrating the same peaks assignment to another homologue series. In the interactive variant of the visualization area selections can be used to evaluate the peak table components contained in the selected part of the plot. In the static plot only the stronger black lines connecting the different peaks of a homologue series are an indicator of overlaps.

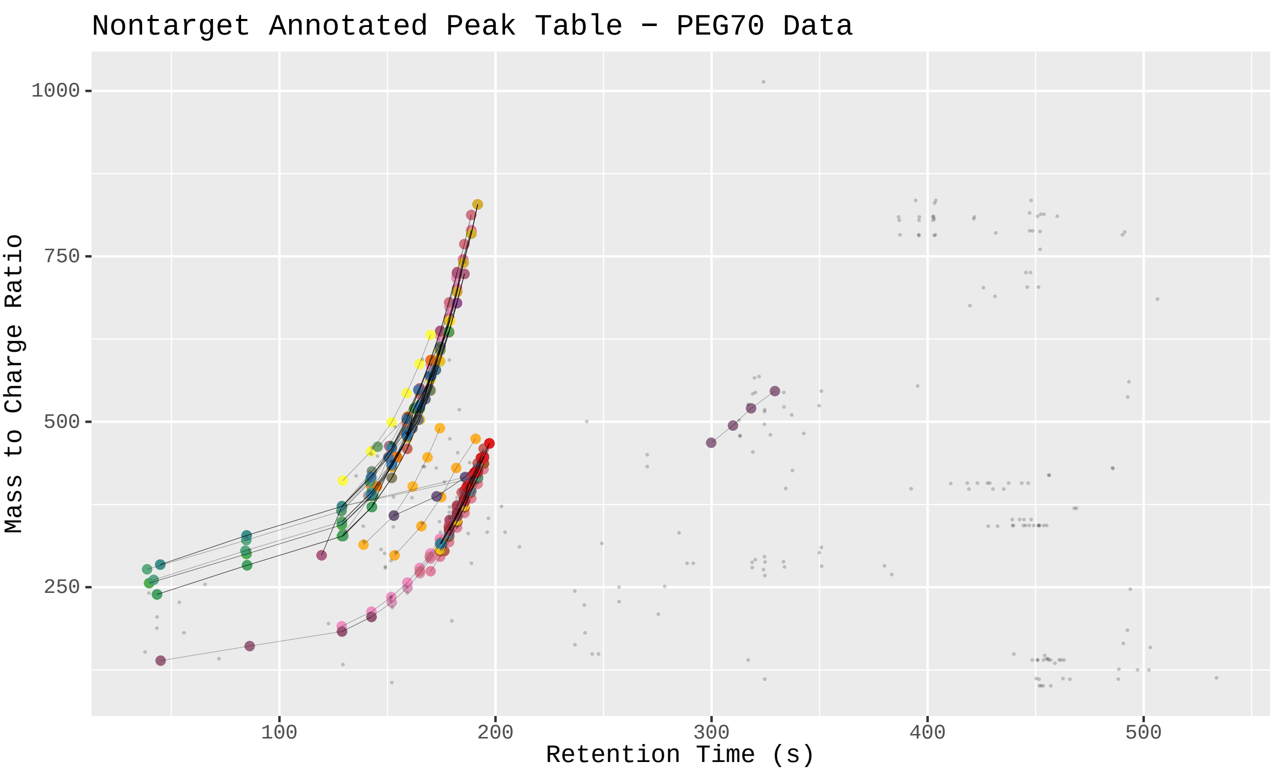

Figure 5: *nontarget* output for plasma samples spiked with PEG70. Peaks not annotated to be part of homologue series are shown as faint gray points. Homologue series peaks are emphasized in size. Members of the same homologue series are connected by a line and share the same color. A custom visualization was implemented to work with *nontarget* output. Peak tables contain duplicate entries for each peak with multiple series assignments. Any peaks highlighted in size that represent homologue series members may thus be overlapping with other points illustrating the same peaks assignment to another homologue series. In the interactive variant of the visualization area selections can be used to evaluate the peak table components contained in the selected part of the plot. In the static plot only the stronger black lines connecting the different peaks of a homologue series are an indicator of overlaps.

#### 2.2 PEG Data Annotation Comparisons

Comparing outputs of *homologueDiscoverer* and *nontarget* shows that both tools agree largely on which peaks are annotated to belong to a homologue series, but differ substantially in exact series of peaks detected. In both PEG17 and PEG70 spikes sample datasets *nontarget* (NT) finds slightly more peaks than *homologueDiscoverer* (HD), with *homologueDiscoverer* annotating 122 of 153 and 150 of 219 peaks to be part of homologue series for the PEG17 and PEG70 datasets respectively (Figures 6 and 7). Exact series overlap between both tools is poorer, with *homologueDiscoverer* agreeing on 14 series with *nontarget* in the PEG17 dataset. However, 5 and 36 series were unique to either *homologueDiscoverer* or *nontarget* (Figure 8). In the PEG70 dataset, differences were even more pronounced, with 15 series being identical across both tools, yet 11 and 96 series being unique to either *homologueDiscoverer* or *nontarget* (Figure 9).

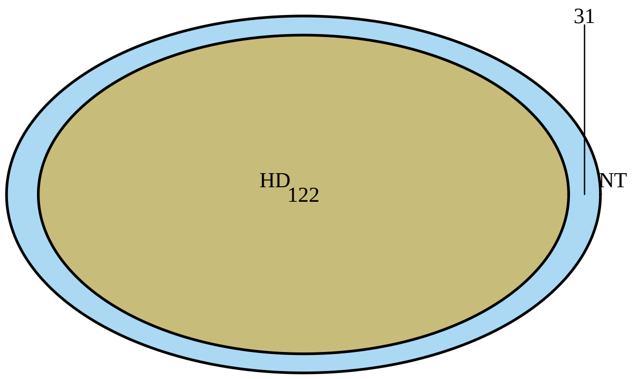

Figure 6: Venn diagram highlighting overlap in annotated peaks between *homologueDiscoverer* (HD) and *nontarget* (NT) for PEG17 dataset.

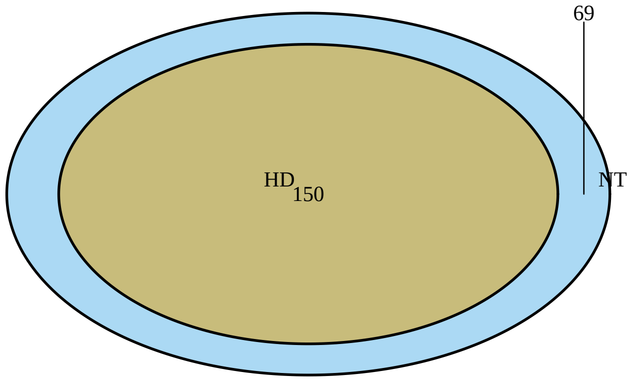

Figure 7: Venn diagram highlighting overlap in annotated peaks between *homologueDiscoverer* (HD) and *nontarget* (NT) for PEG70 dataset.

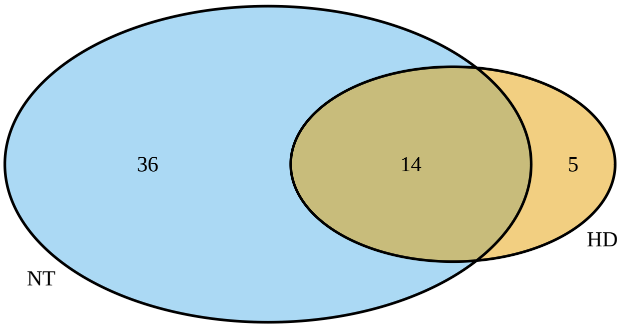

Figure 8: Venn diagram highlighting exact series overlap between *homologueDiscoverer* (HD) and *nontarget* (NT) for PEG17 dataset.

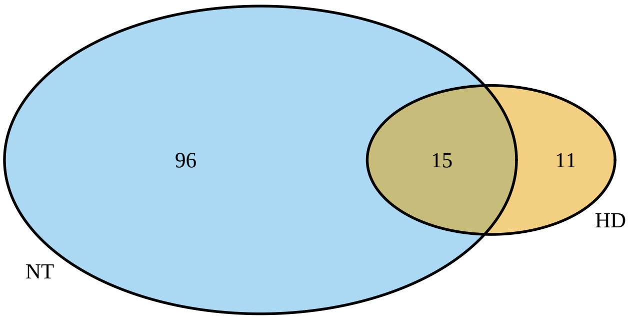

Figure 9: Venn diagram highlighting exact series overlap between *homologueDiscoverer* (HD) and *nontarget* (NT) for PEG70 dataset.

##### 2.3 PEG Data Annotation Validations

For the PEG17 and PEG70 spiked data it is known which peaks originate from the polymer mix and which peaks originate from the plasma sample. While homologue series may also exist in blood plasma samples, we expect more peaks to belong to series in the polymer mix. We do not expect all peaks to belong to homologue series since not all homologue polymers will conform to the heuristic trends we look for.

In the PEG17 dataset *homologueDiscoverer* identifies 115 out of 276 PEG17 mix derived peaks to be part of homologue series, while discovering 7 peaks from the plasma sample to be part of homologue series 10. Results for *nontarget* on this dataset were qualitatively similar, yet with more of the PEG and plasma derived peaks being annotated to be part of homologue series. Here, 143 out of 276 PEG17 derived features were annotated to be part of homologue series, as were 10 plasma derived peaks 11. Results for the PEG70 spiked samples were similar. *homologueDiscoverer* detected 140 out of 251 PEG70 derived peaks to be part of homologue series. 10 plasma peaks were identifier to be part of homologue series. Again, *nontarget* results were more liberal, with 191 of 251 PEG70 derived peaks to be part of homologue series, as were 28 plasma derived peaks.*nontarget* homologue detection results in many peaks which do not belong uniquely to a single homologue series, but rather that many peaks ambiguously belong to many different homologue series (Figure 14).

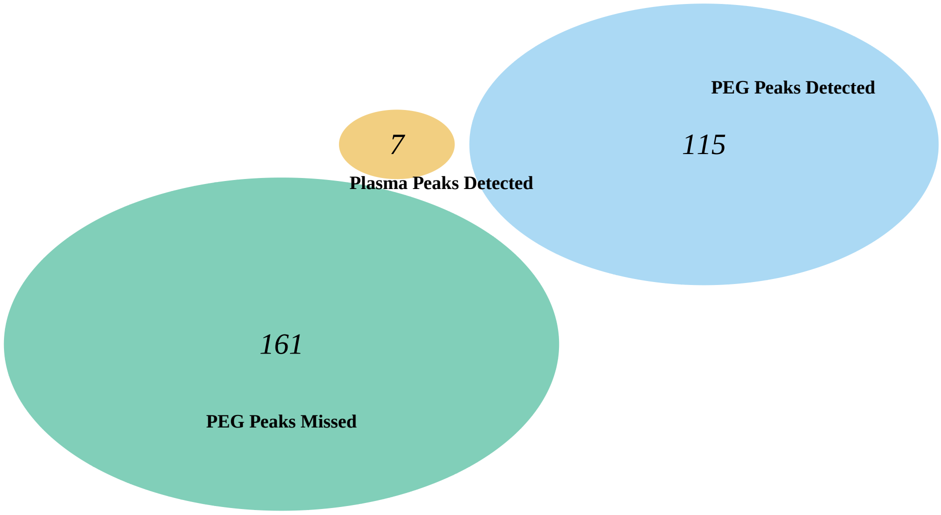

Figure 10: Graphical representation of PEG17 detection performance by *homologueDiscoverer*. Illustrated are peaks matched with the pure PEG samples that were correctly identified to be part of homologue series in mixed plasma and PEG samples, as well as those missed. Plasma derived peaks annotated to be part of homologue series are also highlighted.

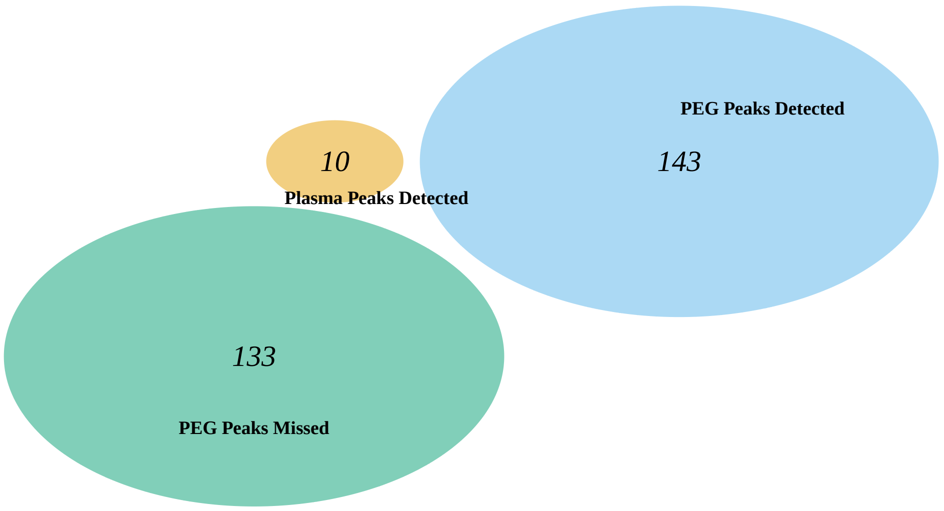

Figure 11: Graphical representation of PEG17 detection performance by *nontarget*. Illustrated are peaks matched with the pure PEG samples that were correctly identified to be part of homologue series in mixed plasma and PEG samples, as well as those missed. Plasma derived peaks annotated to be part of homologue series are also highlighted.

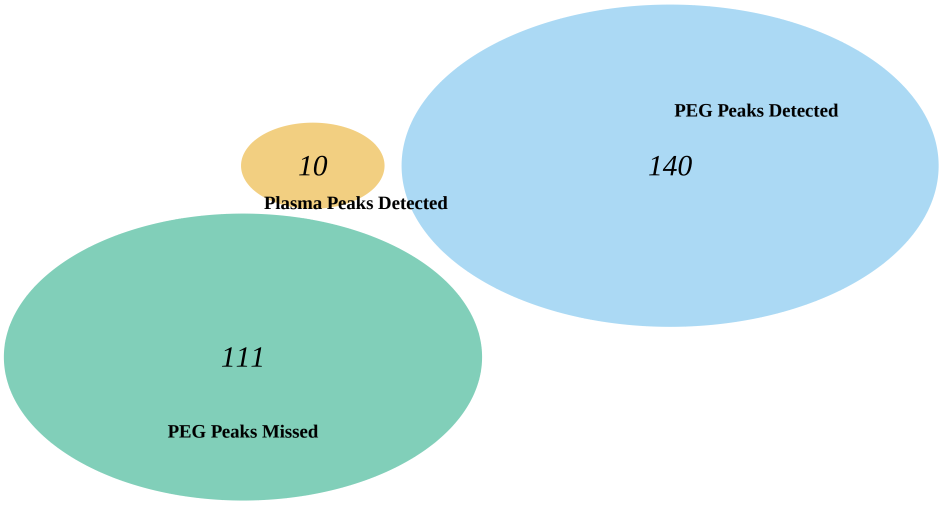

Figure 12: Graphical representation of PEG70 detection performance by *homologueDiscoverer*. Illustrated are peaks matched with the pure PEG samples that were correctly identified to be part of homologue series in mixed plasma and PEG samples, as well as those missed. Plasma derived peaks annotated to be part of homologue series are also highlighted.

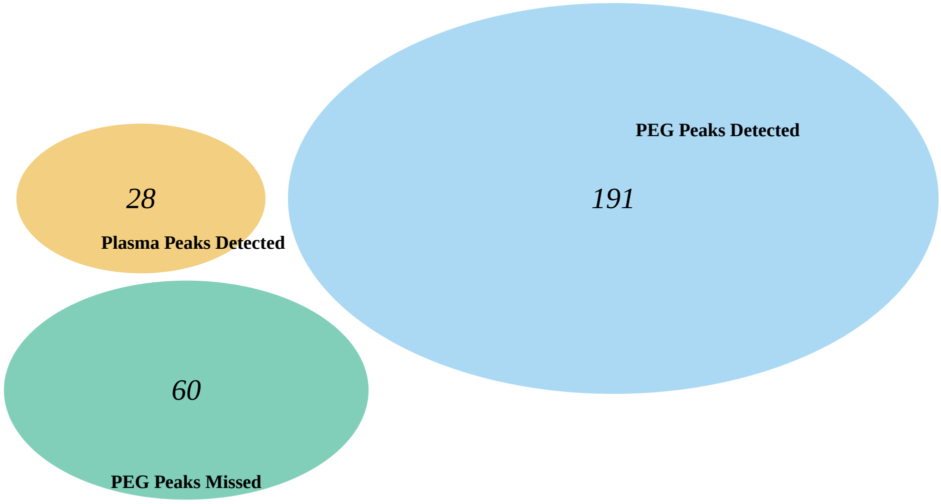

Figure 13: Graphical representation of PEG70 detection performance by *nontarget*. Illustrated are peaks matched with the pure PEG samples that were correctly identified to be part of homologue series in mixed plasma and PEG samples, as well as those missed. Plasma derived peaks annotated to be part of homologue series are also highlighted.

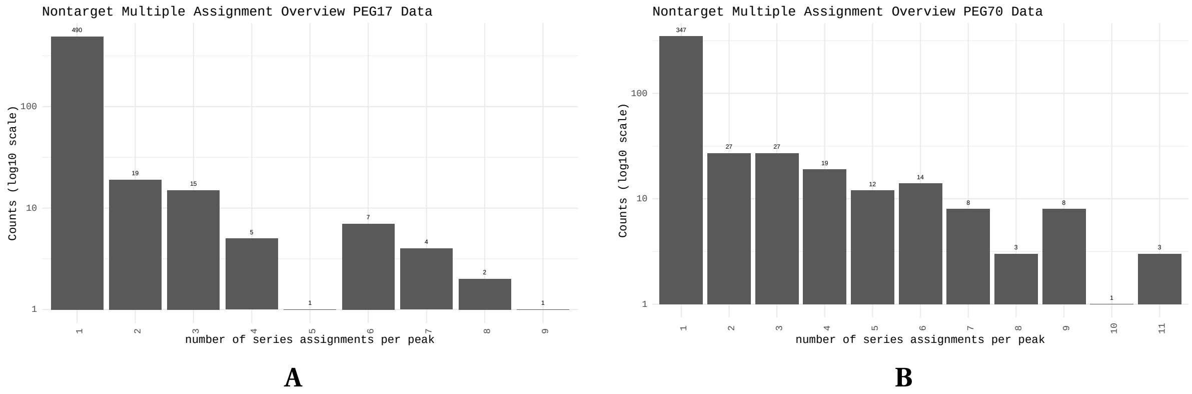

Figure 14: Bar graphs showing the number of multiple homologue series assignments per peak (x-axis) and the frequency of this number of assignments on the y-axis produced by *nontarget* on the PEG17 (**A**) and PEG70 (**B**) datasets.

#### 2.4 PEG Data Results Discussion

*homologueDiscoverer* and *nontarget* both succeed in finding many homologue series in both PEG datasets. Agreement in peaks annotated to be part of homologue series is large, while agreement in exact series is poor. *homologueDiscoverer* uses restrictive heuristics (monotonically increasing or decreasing retention time over mass-to-charge ratio) to exclude possibly spurious series from being returned. *nontarget* makes use of adjusted R2 goodness of fit metrics to exclude series with poor trend consistency. An additional difference between tools is the use of a greedy algorithm in *homologueDiscoverer*, which guarantees that each peak belongs to only one homologue series. The combined effect of forced uniqueness and more restrictive homologue series definition make the output of *homologueDiscoverer* more conservative than the output of *nontarget* which may contain a large number of series assignments per peak.

### 3 MTBLS1358 Data Comparisons

We further compared both *homologueDiscoverer* and *nontarget* on the MTBLS1358 dataset (details in MTBLS1358 data description sub-section here below). *homologueDiscoverer* runs used the untargeted, increment modes with search windows of 13 to 15 for the mass-to-charge ratio and 0.1 to 100 seconds for retention time. Minimum series length was set to 4, and tolerance in ppm was set to 10. Multiple *nontarget* runs were used for comparison purposes. All search window settings were identical, and all other settings left at defaults, except for rttol. rttol was set to 50 for the most comparable run to *homologueDiscoverer*, as well as 0.5 (default value), 1 and 5 for assessing the impact of this setting on *nontarget* output. The code used to generate figures is available on github
https://github.com/kevinmildau/homologueDiscoverer-validation-and-comparison.

#### 3.1 MTBLS1358 Data Description

The MTBLS1358 dataset consists of LC-HRMS measurements of human cell models (HT29 & HepG2), which were treated with a common food contaminant (the mycotoxin deoxynivalenol) to investigate its metabolism in human cells. LC-HRMS analysis was carried out with an QExactive HF instrument coupled to a Vanquish UHPLC system. The raw data was processed with XCMS/MSDial to detect all chromatographic peaks in an untargeted manner. For further information about the dataset the interested readers are referred to the metabolites repository (accession number MTBLS1358) and the publication of Flasch et al. [3]. Data was processed with MS-Dial (version 4.7) [4] in an untargeted fashion to detect all chromatographic peaks in the dataset. Data processing parameters were MS1 tolerance 0.01 Da, Maximum charged number 2, Minimum peak height 1E4 amplitude, Mass slice width 0.01 Da, Smoothing method 'Linear weighted moving average', Smoothing level 3, Minimum peak width 5 scans, Alignment Retention time tolerance 0.05 min, Alignment MS 1 tolerance 0.015 Da. The detected chromatographic peaks were exported to a TSV file for further processing with *homologueDiscoverer*.

#### 3.2 MTBLS1358 Data Annotated Peak Tables

Annotated peak tables of *homologueDiscoverer* (Figure 15) and *nontarget* (Figure 16) qualitatively show the large number of homologue series detected by both tools. We note that the unique assignment of one peak to one homologue series renders the output of *homologueDiscoverer* more readable.

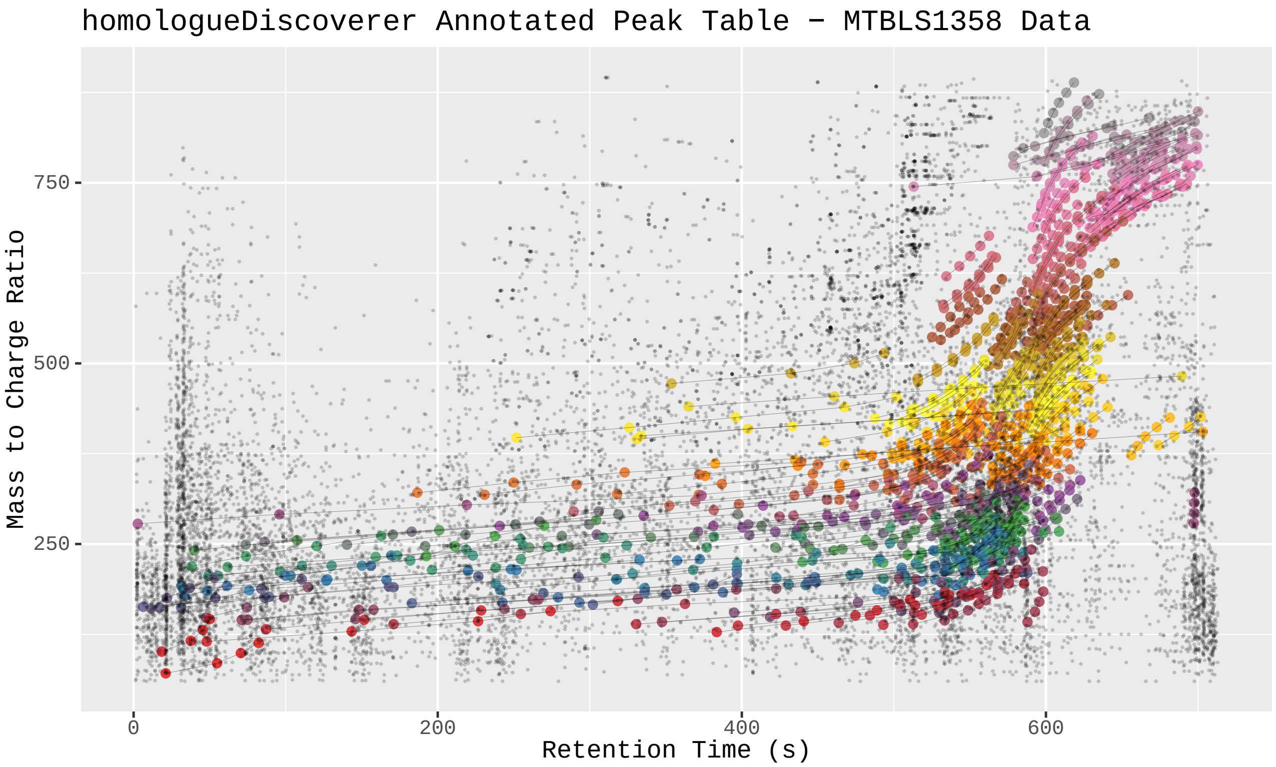

Figure 15: *homologueDiscoverer* output for MTBLS1358 run. Peaks not annotated to be part of homologue series are shown as faint gray points. Homologue series peaks are emphasized in size. Members of the same homologue series are connected by a line and share the same color.

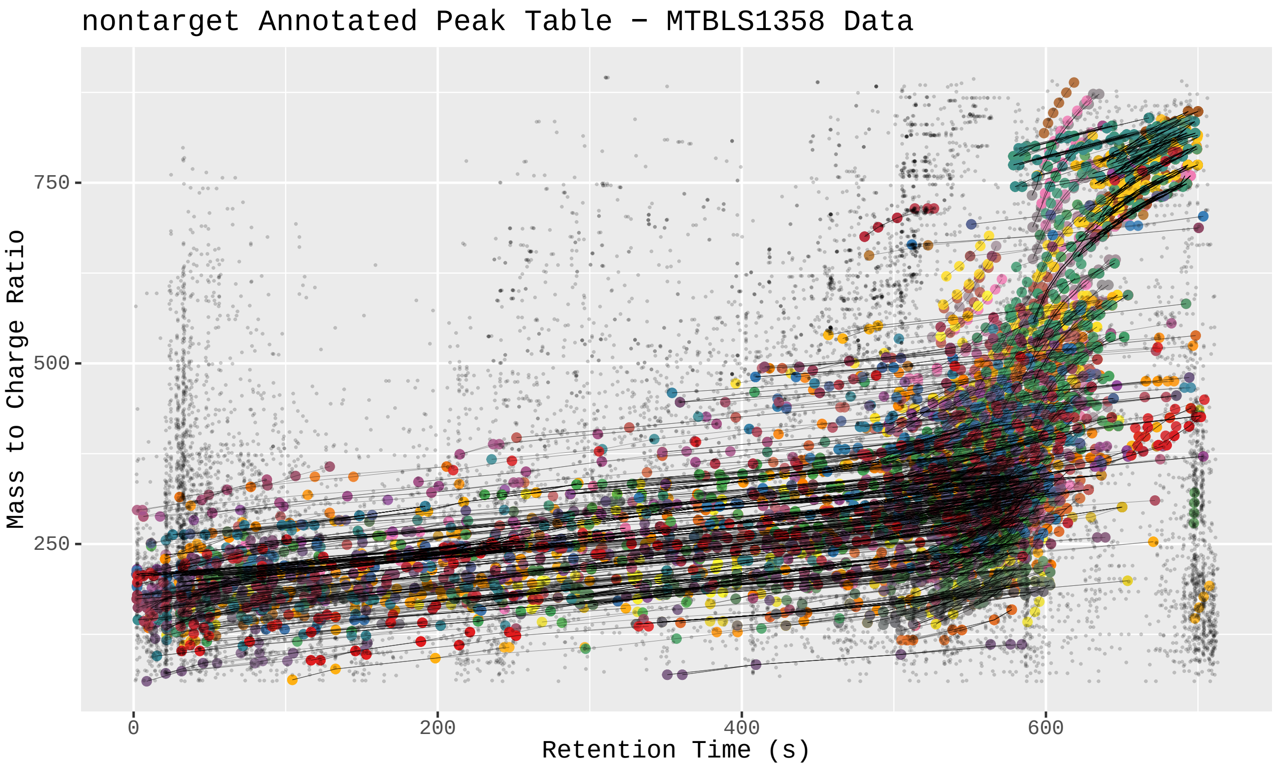

Figure 16: *nontarget* output for MTBLS1358. Peaks not annotated to be part of homologue series are shown as faint gray points. Homologue series peaks are emphasized in size. Members of the same homologue series are connected by a line and share the same color. A custom visualization was implemented to work with *nontarget* output. Peak tables contain duplicate entries for each peak with multiple series assignments. Any peaks highlighted in size that represent homologue series members may thus be overlapping with other points illustrating the same peaks assignment to another homologue series. In the interactive variant of the visualization area selections can be used to evaluate the peak table components contained in the selected part of the plot. In the static plot only the stronger black lines connecting the different peaks of a homologue series are an indicator of overlaps.

##### 3.3 MTBLS1358 Data Annotation Comparisons

Qualitative differences in the number of annotated peaks and found series from the annotated peak tables can be assessed in more detail using the overlap in peaks annotated to be part of homologue series and exact series overlap Venn diagrams in figures 17 and 18. First, we note that the agreement in which peaks are annotated to belong to homologue series is much greater between tools and different runs of *nontarget* than the exact series that are found. Indeed, while 273 peaks are shared in terms of annotated peak overlap, only 6 series are shared as exactly equivalent, i.e., same peak s in same order without any additions or losses, across tools and settings. Furthermore, while annotated peaks between more restricted runs of *nontarget* are largely encompassed by more lenient rttol runs, the same cannot be said for exact series overlap. Indeed, *homologueDiscoverer* finds 159 series unique to itself, while each setting of *nontarget* has a different number of series unique to itself. Most notable is the run with rttol of 50, where 2238 series are found which are not shared with the other runs or *homologueDiscoverer*. Overlap between *homologueDiscoverer* is closest with more lenient rttol settings, as indicated by rttol 50 and rttol 5 runs sharing the largest number of overlaps in both annotated and complete series overlap. Increasing numbers of series coincides with multiple series assignments per peak, where lenient rttol settings lead to many peaks being assigned to belong to numerous homologue series (Figure 19).

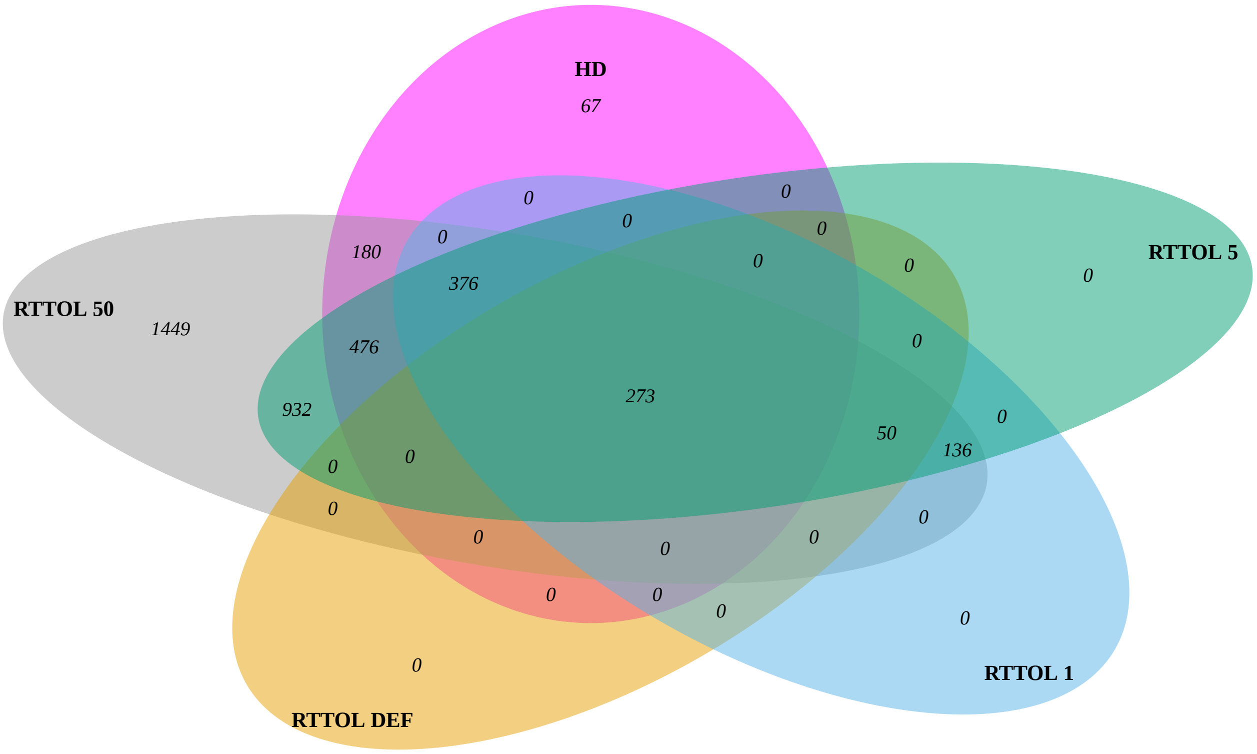

Figure 17: Venn diagram highlighting the overlap in peaks annotated to be part of homologue series between *homologueDiscoverer* and *nontarget* for MTBLS1358 dataset. HD stands for *homologueDiscoverer*, all other homologue search runs were done using *nontarget* and different settings of retention time tolerance (rttol). RTTOL 50 used a tolerance of 50, RTTOL 5 a tolerance of 5, RTTOL 1 a tolerance of 1, and RTTOL DEF used the default tolerance of 0.5.

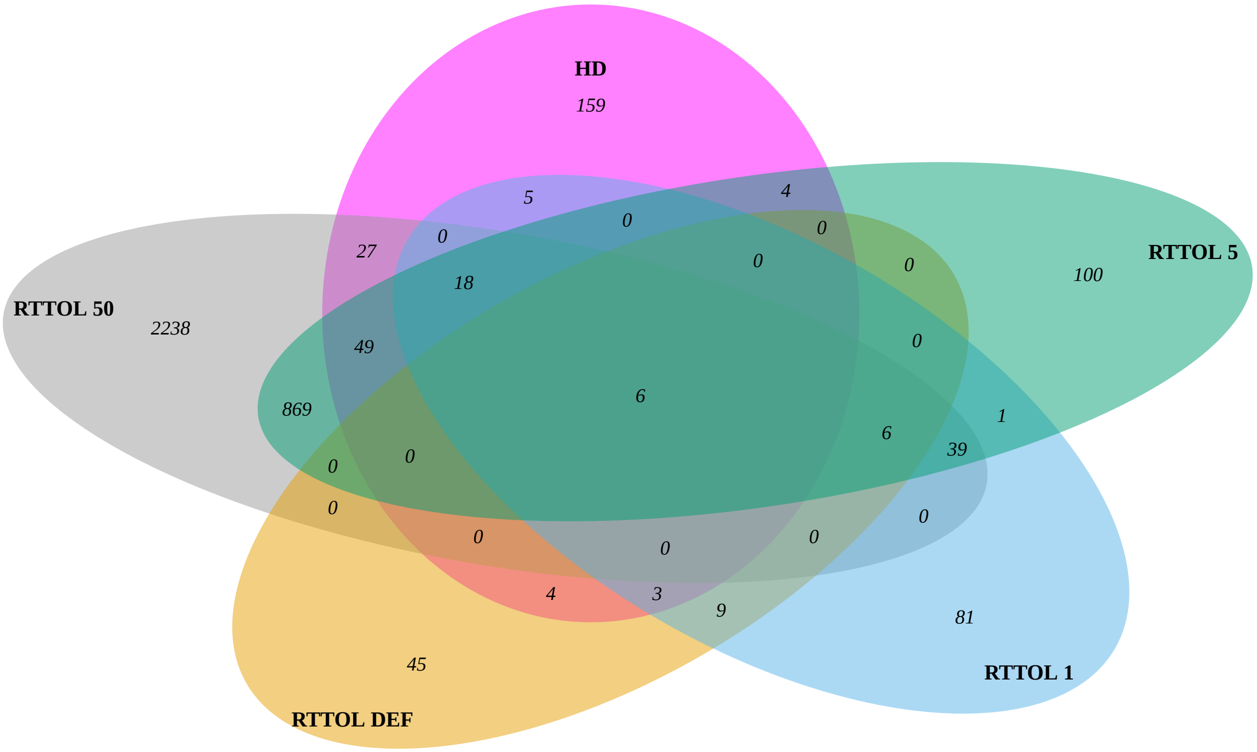

Figure 18: Venn diagram highlighting exact series overlap between *homologueDiscoverer* and *nontarget* for MTBLS1358 dataset. HD stands for *homologueDiscoverer*, all other homologue search runs were done using *nontarget* and different settings of retention time tolerance (rttol). RTTOL 50 used a tolerance of 50, RTTOL 5 a tolerance of 5, RTTOL 1 a tolerance of 1, and RTTOL DEF used the default tolerance of 0.5.

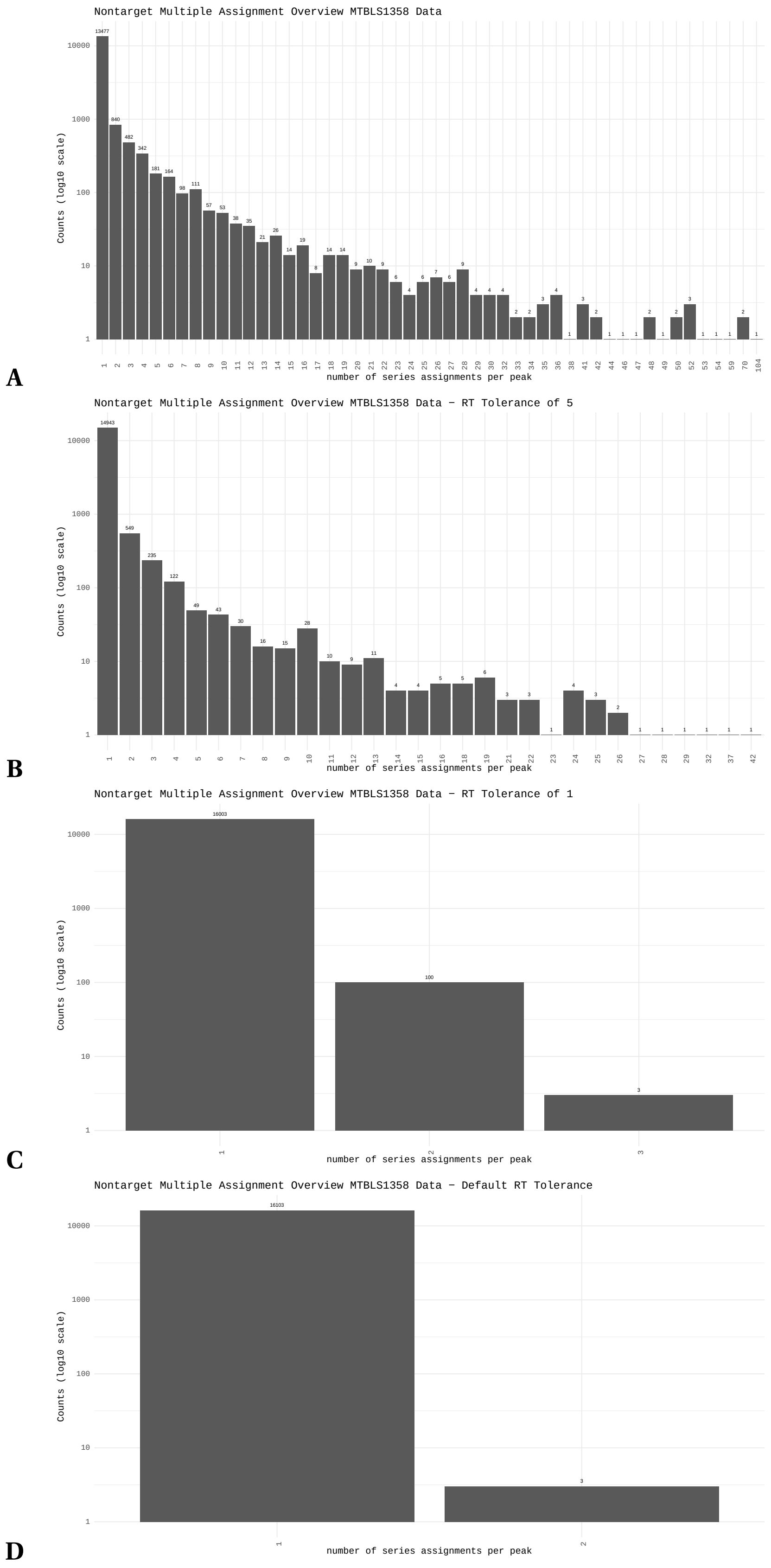

Figure 19: Bar graphs showing the number of multiple homologue series assignments per peak (x-axis) and the frequency of this number of assignments on the y-axis produced by *nontarget* on the MTBLS1358 dataset using different rttol settings. (**A**) Multiple assignment bar graph for rttol of 50. This setting is most comparable to the *homologueDiscoverer* settings used. (**B**) Multiple assignment bar graph for rttol of 5. (**C**) Multiple assignment bar graph for rttol of 1. (**D**) Multiple assignment bar graph for rttol of 0.5. This is the default setting of *nontarget*.

##### 3.4 MTBLS1358 Data Results Discussion

Results obtained for the MTBLS1358 dataset are largely in line with results obtained with the PEG17 and PEG70. However, multiple assignment characteristics of lenient rttol runs of *nontarget* are more pronounced. The large number of multiple assignments leads to almost as many series as there are peaks, rendering the homologue series grouping ineffective at providing a more concise representation of the features. The groupings provided by *homologueDiscoverer* are more concise and we argue that they lend themselves better to manual curation efforts.

##### 3.5 Example Database Output using subset of MTBLS1358 dataset

*homologueDiscoverer* allows storing and summarizing of annotation runs in simple "augmented" peak tables, that is peak tables with additional homologue series information columns. Basic functionality and use cases are illustrated in the package vignette. Here, the vignette output created using the MTBLS1358 dataset provides a simple example of a slice of a series database as shown in Figure 20, and the corresponding series database summary output as shown in Figure 21.

```
# A tibble: 297 × 13
      mz    rt homologue_id within_series_id noise homologue_series_name timestamp
   <dbl> <dbl>        <int>            <int> <lgl> <chr>                 <dttm>
 1  501.  599.            1                1 TRUE  NA                    2022-07-01 12:02:20
 2  515.  603.            1                2 TRUE  NA                    2022-07-01 12:02:20
 3  529.  608.            1                3 TRUE  NA                    2022-07-01 12:02:20
 4  543.  614.            1                4 TRUE  NA                    2022-07-01 12:02:20
 5  557.  623.            1                5 TRUE  NA                    2022-07-01 12:02:20
 6  502.  539.            2                1 TRUE  NA                    2022-07-01 12:02:20
 7  516.  547.            2                2 TRUE  NA                    2022-07-01 12:02:20
 8  530.  555.            2                3 TRUE  NA                    2022-07-01 12:02:20
 9  544.  561.            2                4 TRUE  NA                    2022-07-01 12:02:20
10  558.  566.            2                5 TRUE  NA                    2022-07-01 12:02:20
   push_id sample_peak_id sample_origin sample_description max_intensity normalized_intensity
     <int>          <int> <chr>         <chr>              <lgl>                        <dbl>
 1       1          12408 mtbls1358     NA                 FALSE                       0.0791
 2       1          12635 mtbls1358     NA                 FALSE                       0.215
 3       1          12865 mtbls1358     NA                 TRUE                        0.361
 4       1          13084 mtbls1358     NA                 FALSE                       0.229
 5       1          13275 mtbls1358     NA                 FALSE                       0.116
 6       1          12420 mtbls1358     NA                 TRUE                        0.362
 7       1          12656 mtbls1358     NA                 FALSE                       0.341
 8       1          12877 mtbls1358     NA                 FALSE                       0.139
 9       1          13094 mtbls1358     NA                 FALSE                       0.0929
10       1          13284 mtbls1358     NA                 FALSE                       0.0647
# … with 287 more rows
```

Figure 20: Example of series augmented peak table using a subset of the MTBLS1358 dataset.

```
# A tibble: 49 × 6
   homologue_id series_length  mean_diff median_diff molecular_formula theoretical_increment
          <int>         <int>      <dbl>       <dbl> <chr>                             <dbl>
 1            1             5 14.01571    14.015705  CH2                         14.01565007
 2            2             5 14.01564750 14.01564000 CH2                        14.01565007
 3            3             5 14.01561750 14.01568500 CH2                        14.01565007
 4            4             6 14.015546   14.01556000 CH2                        14.01565007
 5            5             6 14.0156     14.01568000 CH2                        14.01565007
 6            6            10 14.01559778 14.01557000 CH2                        14.01565007
 7            7             6 14.015564   14.01538000 CH2                        14.01565007
 8            8             5 14.01573250 14.01574500 CH2                        14.01565007
 9            9             5 14.01518250 14.015355   CH2                        14.01565007
10           10             8 14.01563286 14.01568000 CH2                        14.01565007
```

Figure 21: Example summary of series augmented peak table using a subset of the MTBLS1358 dataset.

### 4 Yeast Data Comparisons

*homologueDiscoverer* was run on the yeast dataset to illustrate its capacity to find decrementing series (details in yeast data description sub-section here below). Two *homologueDiscoverer* runs were performed with untargeted increment and decrement mode respectively. Search settings included a mass-to-charge ratio window of 10 to 50, and retention time windows of 0.1 to 100 seconds. Minimum series length was set to 6 and tolerance in ppm was to 5. Output from both runs was combined using series database functionalities. A comparable decrement run was performed in *nontarget* using retention time windows of -100 to 0.1 seconds. All non-window settings were left at defaults, expect for rttol which was set at 50 for comparability with *homologueDiscoverer* default settings. The code used to generate figures is available on github `https://github.com/kevinmildau/homologueDiscoverer-validation-and-comparison`.

#### 4.1 Yeast Data Description

The yeast dataset was acquired from ethanolic Pichia pastoris extracts, prepared as described elsewhere [5]. The analysis was conducted on a Thermo Vanquish liquid chromatography system (mounted with a Waters HSS T3 C18 column (2.1 x 150 mm, 1.8 $\mu$m)) coupled to a Q-Exactive-HF orbitrap mass spectrometer. The gradient was composed of two LC-MS grade eluents, eluent A (H2O + 0.1% formic acid) and eluent B (acetonitrile) at 40 degrees Celsius and a flow rate of 0.25 mL/min. After injection of 5 $\mu$L sample, eluent B was kept at 0% for 2 min, then raised to 95% until 13 min, kept at 95% until 15 min, lowered to 0% B, and then kept at 0% until 18 min. The acquired raw files were centroided and converted into mzXML files via ProteoWizard [6]. Data was processed with MS-Dial (version 4.7) [4] in an untargeted fashion to detect all chromatographic peaks in the dataset. Data processing parameters were MS1 tolerance 0.01 Da, Maximum charged number 2, Minimum peak height 1E4 amplitude, Mass slice width 0.01 Da, Smoothing method 'Linear weighted moving average', Smoothing level 3, Minimum peak width 5 scans, Alignment Retention time tolerance 0.05 min, Alignment MS 1 tolerance 0.015 Da. The detected chromatographic peaks were exported to a TSV file for further processing with *homologueDiscoverer*. Since the resulting peak table was very large, all peaks below the third quarter of intensity across all peaks were filtered from the dataset for quicker processing and illustration below.

##### 4.2 Yeast Data Annotated Peak Tables

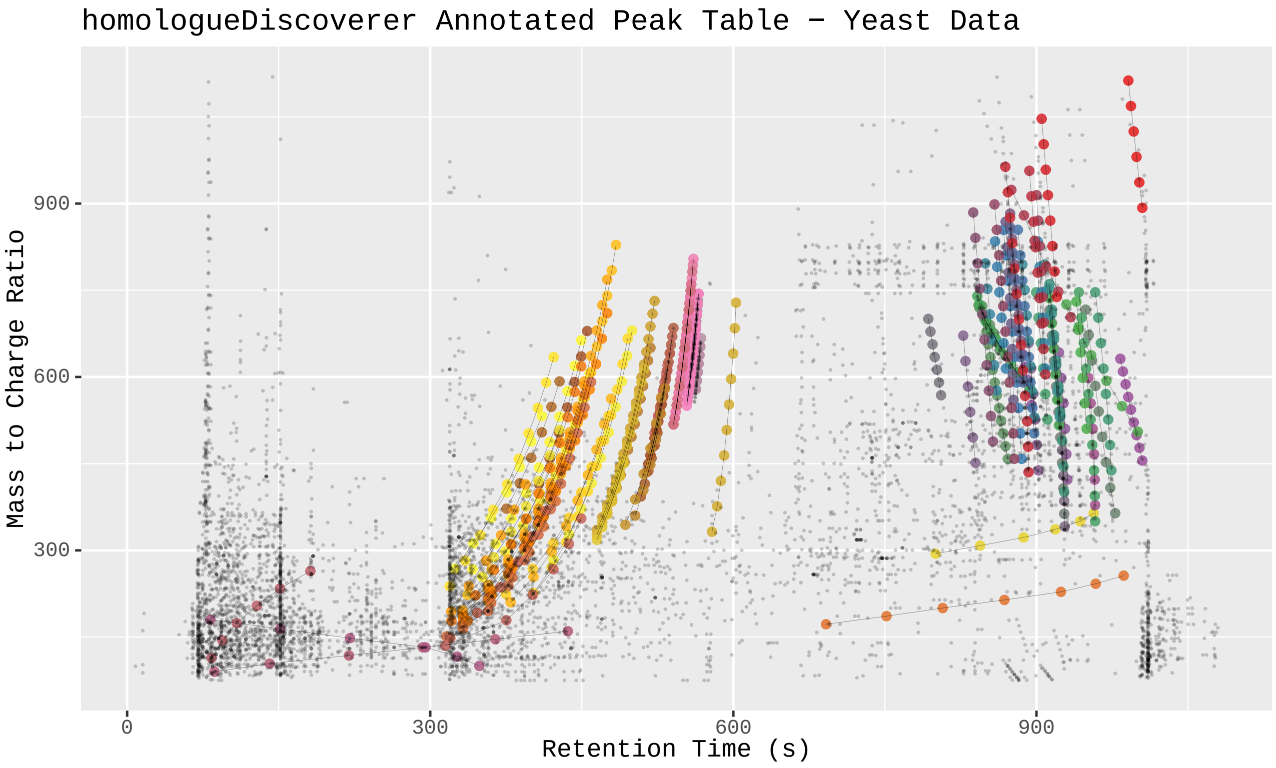

Figure 22: *homologueDiscoverer* output for yeast data. Peaks not annotated to be part of homologue series are shown as faint gray points. Homologue series peaks are emphasized in size. Members of the same homologue series are connected by a line and share the same color. Incrementing and decrementing series were combined.

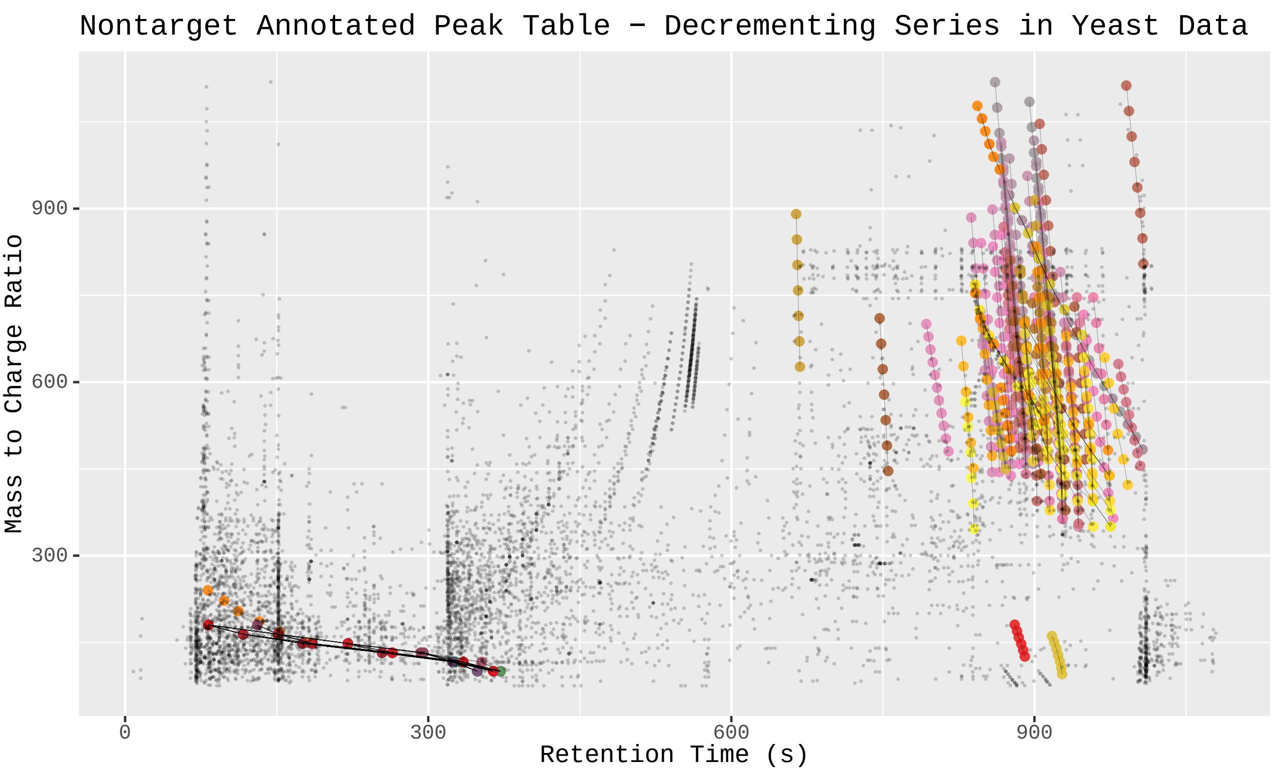

Figure 23: *nontarget* output for yeast data. Peaks not annotated to be part of homologue series are shown as faint gray points. Homologue series peaks are emphasized in size. Members of the same homologue series are connected by a line and share the same color. A custom visualization was implemented to work with *nontarget* output. Peak tables contain duplicate entries for each peak with multiple series assignments. Any peaks highlighted in size that represent homologue series members may thus be overlapping with other points illustrating the same peaks assignment to another homologue series. In the interactive variant of the visualization area selections can be used to evaluate the peak table components contained in the selected part of the plot. In the static plot only the stronger black lines connecting the different peaks of a homologue series are an indicator of overlaps. Only decrementing series are shown since combining output from multiple runs was non-trivial.

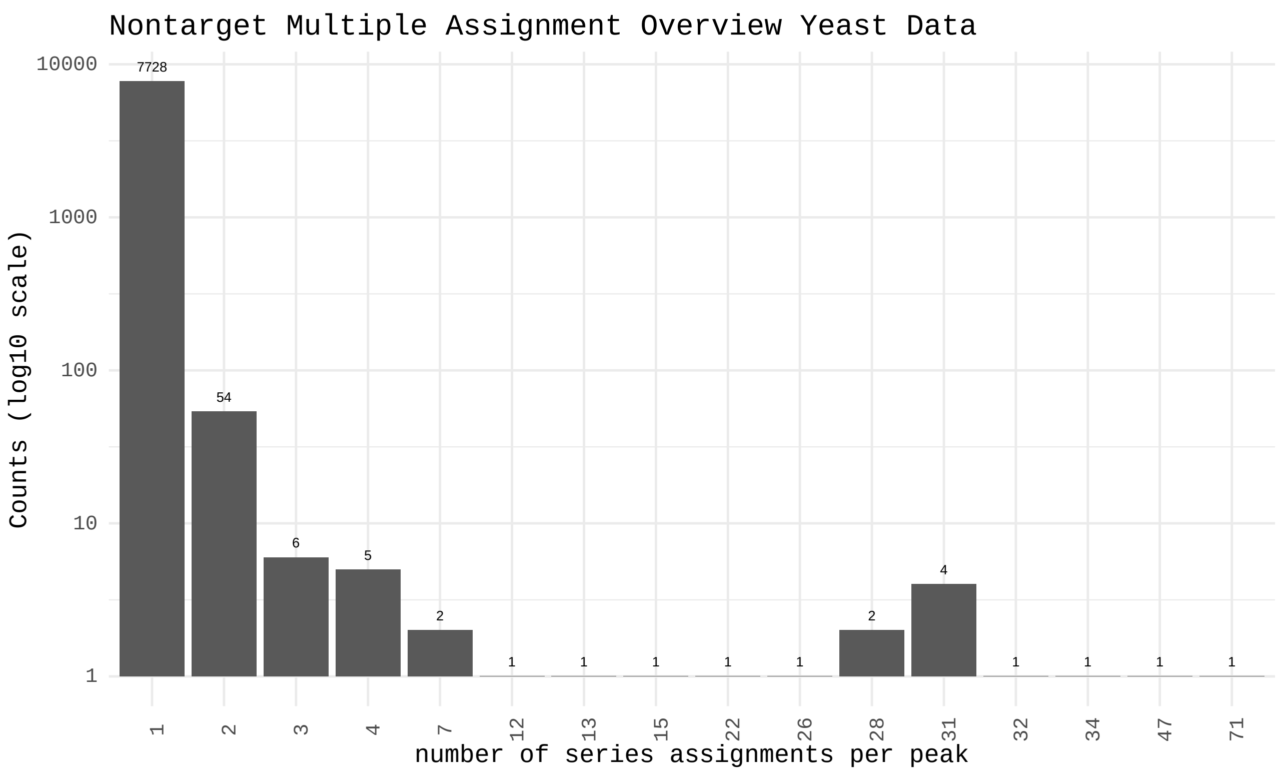

Figure 24: Bar graph showing the number of multiple homologue series assignments per peak (x-axis) and the frequency of this number of assignments on the y-axis produced by *nontarget* on the yeast dataset.

##### 4.3 Yeast Data Results Discussion

The yeast dataset illustrates the presence of increasing and decreasing mass-to-charge ratio over retention time trends. Both *homologueDiscoverer* and *nontarget* can be used to search for these trends (Figures 22 and 23). Using the database functionality implemented in *homologueDiscoverer*, the results of both runs are easily merged for full results visualization (Figure 22). As was observed with the previous datasets, *nontarget* produces many series assignments per peak for the yeast dataset as well (Figure 24).
